## Supplementary Material for "Alterations in cortical excitability during pain: A combined TMS-EEG Study"

^1^Center for Pain IMPACT, Neuroscience Research Australia, Sydney, New South Wales, Australia, ^2^University of New South Wales, Sydney, New South Wales, Australia, ^3^School of Health Sciences, College of Health, Medicine and Wellbeing, The University of Newcastle, Callaghan, New South Wales, Australia, ^4^Department of Neural and Pain Sciences, School of Dentistry, University of Maryland Baltimore, ^5^Center to Advance Chronic Pain Research, University of Maryland Baltimore, ^6^Department of Medical Biophysics, University of Western Ontario, Canada, ^7^The Gray Centre for Mobility and Activity, University of Western Ontario, London, Canada.

**Supplementary Results for Experiment 1**

***Excluded Channels, Epochs and ICA Components***

The mean number of channels removed across participants was 2.2 ± 2.7. The mean number of epochs excluded was 5.8 ± 5.2, 5.1 ± 5.0 and 5.3 ± 4.1 for the pre-pain, pain and post-pain conditions respectively. The mean number of components rejected was 11.0 ± 6.3, 11.7 ± 8.4 and 10.25 ± 8.3 for the pre-pain, pain and post-pain block respectively. A Bayesian one-way ANOVA revealed moderate evidence for no difference in number of rejected epochs (BF_10_ = 0.145), or components (BF_10_ = 0.193), between conditions

***Postpain comparisons***

Figure S1 shows the TEPs across pre-pain vs. post-pain (S1A) and pain vs. post-pain (S1B) conditions for the frontocentral electrodes (left) and parietal-occipital electrodes identified from the cluster analysis. For prepain vs. post-pain, Bayes paired t-tests comparing peak amplitude between conditions yielded anecdotal evidence of no difference for the frontocentral N45 (BF_10_ = 0.465) and the parietal-occipital P60 (BF_10_ = 0.351) and moderate evidence for no difference in the frontocentral N100 (BF_10_ = 0.205). For pre-pain vs. post-pain, Bayes paired t-tests comparing peak amplitude between conditions yielded anecdotal evidence of no difference for the frontocentral N45 (BF_10_ = 0.992), frontocentral N100 (BF_10_ = 0.775), and parietal-occipital P60 (BF_10_ = 0.692).

***Relationships between changes in MEP amplitude and changes in N45, N100, P60 amplitude during pain***

Respectively, there was moderate and anecdotal evidence for no relationship between alterations in MEP amplitude and alterations in the N100 (r_25_ = -.14, BF_10_ = 0.303) and P60 (r_25_ = 0.185, BF_10_ = 0.36) during pain. There was anecdotal evidence for a relationship between alterations in the N45 and MEP amplitude during pain (r_25_ = -.387, BF_10_ = 1.40)

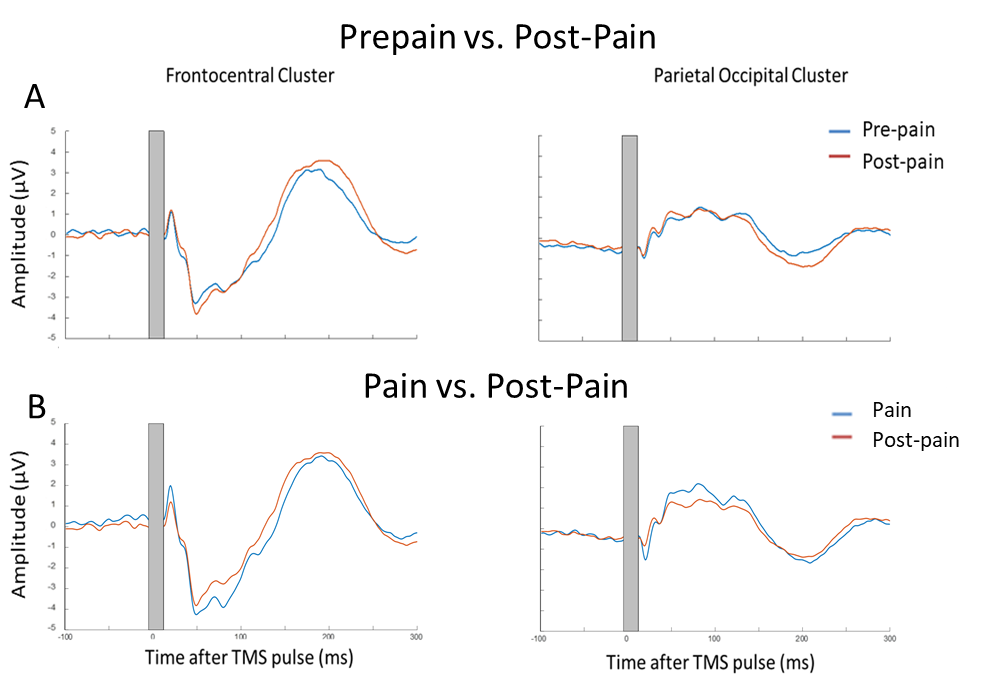

*Figure S1.* TEPs for pre-pain vs. post-pain (Panel A) and pain vs. post-pain (Panel B) conditions and pre for the frontocentral electrodes (left) and parietal-occipital electrodes (right) identified from the cluster analysis. The grey shaded area represents the window of interpolation around the transcranial magnetic stimulation (TMS) pulse.

**Supplementary Results for Experiment 2**

The mean number of channels removed across participants was 3 ± 2.2. The mean number of epochs excluded was 5.1 ± 3.0, 5.6 ± 3.4, 8.2 ± 5.2 and 4.6 ± 4.4 for the active pre-pain, sham pre-pain, active pain, and sham pain conditions respectively. The mean number of ICA components rejected was 11.4 ± 8.4, 12.3 ± 6.9, 10.1 ± 3.4 and 12.9 ± 9.5 for the active pre-pain, sham pre-pain, active pain and sham pain conditions respectively. A Bayesian one-way ANOVA revealed moderate evidence for no difference in number of rejected epochs (BF_10_ = 0.27), or components (BF_10_ = 0.181), between conditions

The mean cold and warm detection threshold was 29.6 ± 2.6°C and 36.0 ± 3.6°C respectively. The mean heat pain threshold was 42.1 ± 3.6 °C.

The mean RMT and test stimulus intensity was, respectively, 70.4 ± 4.3% and 77.7 ± 4.7% of maximum stimulus output. The mean test electrical stimulation intensity for the sham TMS condition was 4.4 ± 2.5 mA. This intensity is comparable to previous studies using sham electrical stimulation and is insufficient to directly activate the cortex (Nahian S Chowdhury, Rogasch, et al., 2022; Conde et al., 2019; Rocchi et al., 2021)

Figure S2 shows the mean ratings during the 6 thermal stimuli delivered during the pain block, for both active and sham TMS. All participants reported 0/10 pain during the pre-pain block.

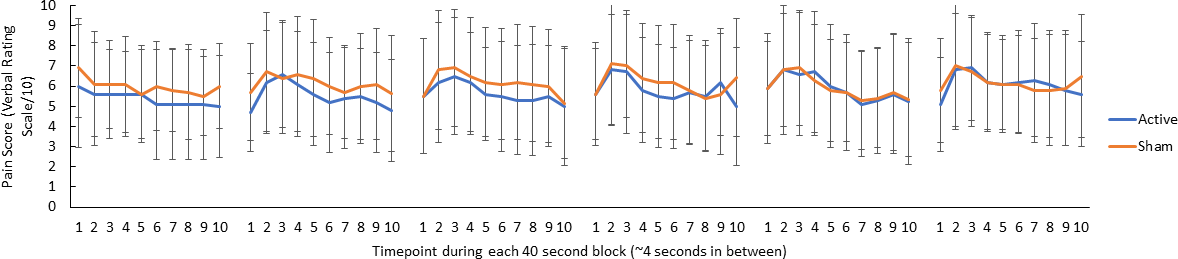

*Figure S2.* Mean (± SD) pain ratings during the 6 thermal stimuli delivered during the pain block (thermal stimuli delivered at 46°C) for both active and sham TMS. 10 pain ratings were collected over each 40 second stimulus ~ every 4 second.

**Supplementary Results for Experiment 3**

The mean number of channels removed across participants was 6.1 ± 2.64. The mean number of epochs excluded was 5.4 ± 3.4, 8.1 ± 6.5, 6.5 ± 6.6 and 5.4 ± 2.8 for the subthreshold pre-pain, suprathreshold pre-pain, subthreshold pain, suprathreshold pain respectively. The mean number of ICA components rejected was 9.1 ± 7.8, 7.6 ± 7.1, 9.6 ± 7.8 and 8.6 ± 7.0 for the subthreshold pre-pain, suprathreshold pre-pain, subthreshold pain, suprathreshold pain. A Bayesian one-way ANOVA revealed moderate evidence for no difference in number of rejected epochs between conditions (BF_10_ = 0.169), and no conclusive evidence for a difference in number of rejected components between conditions (BF_10_ = 0.576).

The mean cold and warm detection threshold was 29.4 ± 2.0°C and 34.9 ± 1.4°C respectively. The mean heat pain threshold was 42.9 ± 2.5°C.

The mean RMT, subthreshold and suprathreshold stimulus intensity was, respectively, 68.0 ± 8.3%, 61.0 ± 7.3 % and 74.8 ± 9.1% of maximum stimulus output.

Figure S3 shows the mean warmth ratings during the 6 thermal stimuli delivered during the pre-pain block, for both supra- and subthreshold TMS, confirming that participants felt some difference in the sensation (relative to baseline) when the temperature was delivered at the warmth detection threshold.

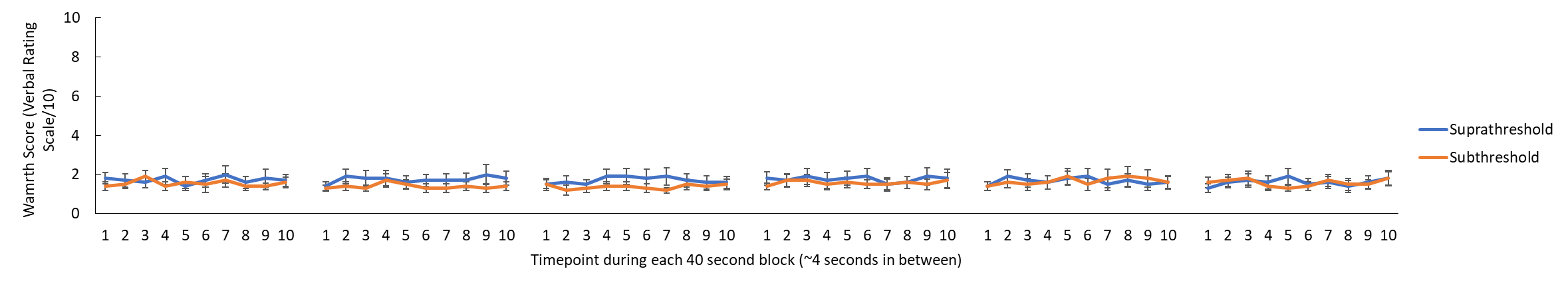

*Figure S3.* Mean (± SEM) warmth ratings during the 6 thermal stimuli delivered during the pre-pain block (thermal stimuli delivered at the warmth threshold) for both supra and subthreshold TMS. 10 warmth ratings were collected over each 40 second stimulus ~ every 4 second.

Figure S4 shows the mean pain ratings during the 6 thermal stimuli delivered during the pain block, for both supra- and subthreshold TMS. All participants reported 0/10 pain during the pre-pain block.

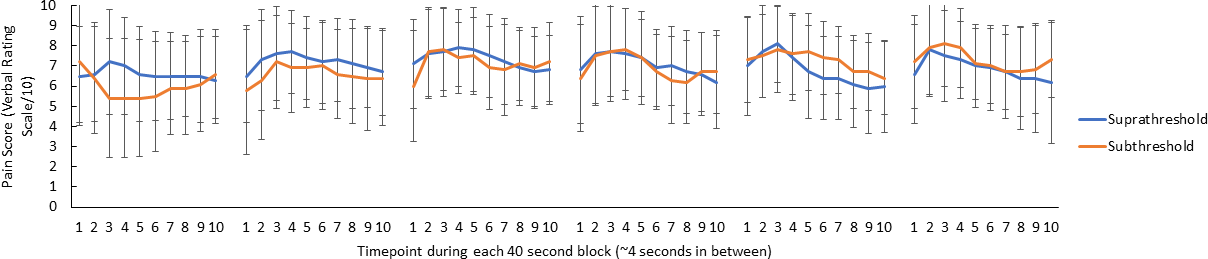

*Figure S4.* Mean (± SD) pain ratings during the 6 thermal stimuli delivered during the pain block (thermal stimuli delivered at 46°C) for both supra and subthreshold TMS. 10 pain ratings were collected over each 40 second thermal stimulus ~ every 4 second.

**Test-retest reliability of N45 Peaks**

We conducted a supplementary investigation of the test-retest reliability of the N45 TEP peaks for each experiment. The interclass correlation coefficient (Two-way fixed, single measure) for the N45 to active suprathreshold TMS across timepoints for each experiment was 0.90 for Experiment 1 (across pre-pain, pain, post-pain time points), 0.74 for Experiment 2 (across pre-pain and pain conditions), and 0.95 for Experiment 3 (across pre-pain conditions). This suggests that even with the fluctuations in the N45 induced by pain, the N45 for each participant was stable across time, further supporting the reliability of our data.
